# The potential for mobile demersal fishing to reduce carbon storage and sequestration in seabed sediments

**DOI:** 10.1101/2021.07.07.450307

**Authors:** Graham Epstein, Julie P. Hawkins, Catrin R. Norris, Callum M. Roberts

## Abstract

Subtidal marine sediments are one of the planet’s primary carbon stores and strongly influence the oceanic sink for atmospheric CO_2_. By far the most pervasive human activity occurring on the seabed is bottom trawling and dredging for fish and shellfish. A global first-order estimate suggested mobile demersal fishing activities may cause 160-400 Mt of organic carbon (OC) to be remineralised annually from seabed sediment carbon stores. There are, however, many uncertainties in this calculation. Here, we discuss the potential drivers of change in seabed OC stores due to mobile demersal fishing activities and conduct a systematic review, synthesising studies where this interaction has been directly investigated. Mobile demersal fishing would be expected to reduce OC in seabed stores, albeit with site-specific variability. Reductions would occur due to lower production of flora and fauna, the loss of fine flocculent material, increased sediment resuspension, mixing and transport, and increased oxygen exposure. This would be offset to some extent by reduced faunal bioturbation and respiration, increased off-shelf transport and increases in primary production from the resuspension of nutrients. Studies which directly investigated the impact of demersal fishing on OC stocks had mixed results. A finding of no significant effect was reported in 51% of 59 experimental contrasts; 41% reported lower OC due to fishing activities, with 8% reporting higher OC. In relation to remineralisation rates within the seabed, 14 experimental contrasts reported that demersal fishing activities decreased remineralisation, with four reporting higher remineralisation rates. The direction of effects was related to sediment type, impact duration, study design and local hydrography. More evidence is urgently needed to accurately quantify the impact of anthropogenic physical disturbance on seabed carbon in different environmental settings, and incorporate full evidence-based carbon considerations into global seabed management.

## 1. Introduction

Through a mixture of physical, chemical and biological processes, the ocean has absorbed ~40% of anthropogenic CO_2_ emissions since the industrial revolution (Gruber et al. 2019, Watson et al. 2020). The term “blue carbon” describes the ability of marine ecosystems to absorb CO_2_ from the atmosphere or water column, assimilate this inorganic carbon (IC) into organic compounds and isolate it from remineralisation for centennial to millennial time-scales (Nellemann et al. 2009). This process of carbon capture is key to maintaining the ecological functioning of the ocean (Bauer et al. 2013) and is beneficial as a sink for anthropogenic CO_2_ (Khatiwala et al. 2009, Gruber et al. 2019).

Research on blue carbon initially focused on the coastal vegetated habitats of mangroves, seagrass and saltmarsh, due to their ability to fix CO_2_ directly, store high concentrations of organic carbon (OC) *in-situ* within underlying sediments and to accrete this OC indefinitely over time (McLeod et al. 2011, Duarte et al. 2013). Although these habitats are some of the most intense OC sinks on the planet (Duarte et al. 2013), with sequestration rates considerably higher than forests on land (McLeod et al. 2011), their limited spatial scale of approximately 1 million km^2^ or ~0.2% of the ocean’s surface, means they only contain a small proportion of the ocean’s total OC stock (Nellemann et al. 2009, Duarte et al. 2013, Duarte 2017, Howard et al. 2017, Atwood et al. 2020).

By far the largest mass of OC occurs in the pelagic zone (Nellemann et al. 2009), with much of this in flux between the oceanic IC pool and the atmosphere (Bauer et al. 2013). However, at depths below 1000 m, pelagic OC may become isolated from atmospheric exchange processes for centennial time scales (Caldeira et al. 2002, Nellemann et al. 2009, Krause-Jensen and Duarte 2016). How this should be accounted for remains a matter of debate, and so pelagic OC is rarely used in the quantification or classification of blue carbon (Lovelock and Duarte 2019). That withstanding, subtidal marine sediments contain the ocean’s biggest OC store, estimated to hold ~87 Gt of OC in the upper 5 cm (Lee et al. 2019) or ~2.3 Tt in the top 1 m (Duarte et al. 2013, Atwood et al. 2020). Quantification of annual sequestration rates in these sediments is relatively poorly constrained, however they have been estimated globally at approximately 126 - 350 Mt OC yr^-1^ (Berner 1982, Seiter et al. 2004, Burdige 2007, Keil 2017, Lee et al. 2019, Smeaton et al. 2021).

Seabed sediments are subjected to a wide range of direct physical impacts from human pressures, namely: shipping, mineral extraction, fishing, energy developments, deployment of cables and pipelines, coastal development, dredging of shipping access channels and disposal of dredge spoil (Halpern et al. 2019, O’Hara et al. 2021). By far the most widespread source of disturbance is bottom trawling and dredging for fish and shellfish (Oberle et al. 2016a, Amoroso et al. 2018, Kroodsma et al. 2018, O’Hara et al. 2021). These pressures are pervasive and long lasting, with improved technologies over the last two centuries increasing the spread of mobile fishing gears to deeper waters and much of the global ocean (Roberts 2007, Watson and Morato 2013, Kroodsma et al. 2018). Compared to many other types of stressors, in intensively fished areas, trawling and dredging can also occur on the same area of seabed numerous times in a year (Tillin et al. 2006, Hinz et al. 2009, Oberle et al. 2016a).

Globally, fishing pressure with mobile demersal gear is concentrated in subtidal areas at depths above 1000 m in coastal habitats and offshore on continental shelves and slopes (Amoroso et al. 2018, Kroodsma et al. 2018). In total these areas cover around 9% of the global seabed, yet they store an estimated 360 Gt in their top 1 m of sediment (Atwood et al. 2020). Continental shelf sediments are also highly productive, estimated to sequester up to 86% of all OC that is buried annually in global subtidal sediments (Berner 1982, Seiter et al. 2004, Atwood et al. 2020, Smeaton et al. 2021).

Mobile demersal fishing activity significantly alters seabed faunal communities (Kaiser et al. 2006, Sciberras et al. 2016, Hiddink et al. 2017), restructures the top layers of benthic sediments (Trimmer et al. 2005, Puig et al. 2012, Eigaard et al. 2016, Oberle et al. 2016b) and resuspends large volumes of sediment into the water column (Jones 1992, Ruffin 1998, Thrush and Dayton 2002, Durrieu de Madron et al. 2005, Martín et al. 2014b, Palanques et al. 2014). However, the net effect of this disturbance on OC stores is poorly resolved. Through mixing, resuspension and oxidation of surface sediments, along with the disturbance of benthic communities, fishing likely generates a source of “underwater carbon emissions” via increased remineralisation of OC, and will also limit future OC sequestration by inhibiting long-term sediment settlement and consolidation (Martín et al. 2014b, Keil 2017, Luisetti et al. 2019, De Borger et al. 2021, Sala et al. 2021). This disturbance can be expected to increase IC concentrations in the ocean, and via this, slow the rate of CO_2_ uptake from the atmosphere, while contributing to ocean acidification and potentially leading to increased release of oceanic CO_2_ to the atmosphere (Khatiwala et al. 2009, Pendleton et al. 2012, Bauer et al. 2013, Keil 2017, Lovelock et al. 2017, Luisetti et al. 2019, LaRowe et al. 2020, Sala et al. 2021). To place the effect of mobile demersal fishing in full context, it is important to better quantify the impacts of different pressures on OC storage and to understand how these compare with natural hydrological disturbances to seabed sediments in different environmental settings (Winterwerp and Kranenburg 2002, Pusceddu et al. 2005b, Arndt et al. 2013, Rühl et al. 2020).

The cycling and storage of OC at the seabed is highly complex and is influenced by: sediment fauna, flora and microbiome; its lithology and granulometry; and the chemistry, hydrology and biology of the surrounding water column (Burdige 2007, Bauer et al. 2013, Keil 2017, Middelburg 2018, Snelgrove et al. 2018, LaRowe et al. 2020, Rühl et al. 2020). With all of these factors affected by many positive and negative feedback mechanisms, it is challenging to definitively identify the impact of trawling and dredging on net OC storage (Keil 2017, Snelgrove et al. 2018, LaRowe et al. 2020, Rühl et al. 2020). In this review we discuss the potential drivers of change in sediment OC due to mobile demersal fishing activities, and summarise empirical evidence where their effects on sediment OC has been investigated. We also discuss recent peer reviewed publications which aim to quantify the impact of mobile demersal fishing at global, regional and national scales, and highlight why the results must be viewed with both concern and caution (Luisetti et al. 2019, Paradis et al. 2021, Sala et al. 2021). If seabed sediments were to be recognised as a quantifiable and manageable blue carbon resource it could unlock huge climate change mitigation potential and carbon financing opportunities (Avelar et al. 2017, Seddon et al. 2019).

## 2. Links between seabed sediment OC and mobile demersal fishing

### 2.1 Production of benthic micro- and macroalgae

Seabed OC is mostly allochthonous, with much of it originating from terrestrial run-off and primary production in surface waters from phytoplankton, macroalgae and wetland vegetation (Bauer et al. 2013, Turner 2015, Krause-Jensen and Duarte 2016, LaRowe et al. 2020, Legge et al. 2020). Through the ocean’s “biological pump” much of this OC will be consumed, repackaged, excreted or remineralised before a remaining proportion of OC reaches the seabed (Turner 2015, Keil 2017, Middelburg 2018). On sediments in the euphotic zone, some OC is autochthonous – i.e. produced *in-situ* by microphytobenthos, and by macroalgae found on more stable sediments, hard substrate or attached to biogenic material (MacIntyre et al. 1996, Gattuso et al. 2006).

While the impact of mobile demersal fishing on benthic algae is little studied, it is known that benthic macroalgae are easily damaged by physical disturbance, and the structure and abundance of microphytobenthos is highly dependent on both natural and anthropogenic perturbation (MacIntyre et al. 1996, Fragkopoulou et al. 2021). In general, mobile demersal fishing is expected to lead to a reduction in algal cover and sediment surface chlorophyll a concentration (Fig. 1a) (Mayer et al. 1991, MacIntyre et al. 1996, Watling et al. 2001, Tiano et al. 2019, Fragkopoulou et al. 2021). For example, scallop dredging at depths of 8-15 m in the Damariscotta River Estuary of the Northwest Atlantic led to clear visual disturbance of diatom matts and caused a significant reduction in chlorophyll a concentration (Mayer et al. 1991, Watling et al. 2001). Among algae, kelp and coralline algae can require years and decades respectively, to recover following disturbance (Dayton et al. 1992, Fragkopoulou et al. 2021). By contrast ephemeral macroalgae and microphytobenthos can recover quickly, especially from less chronic disturbance (MacIntyre et al. 1996, Ordines et al. 2017). For example, in the *Pesquera Rica* trawling grounds of the Balearic Islands, red algae beds of Peyssonneliaceae and Corallinophycidae persist within trawled areas, although their biomass is around 39-47% lower compared to untrawled areas (Ordines et al. 2017).

**Figure 1.**
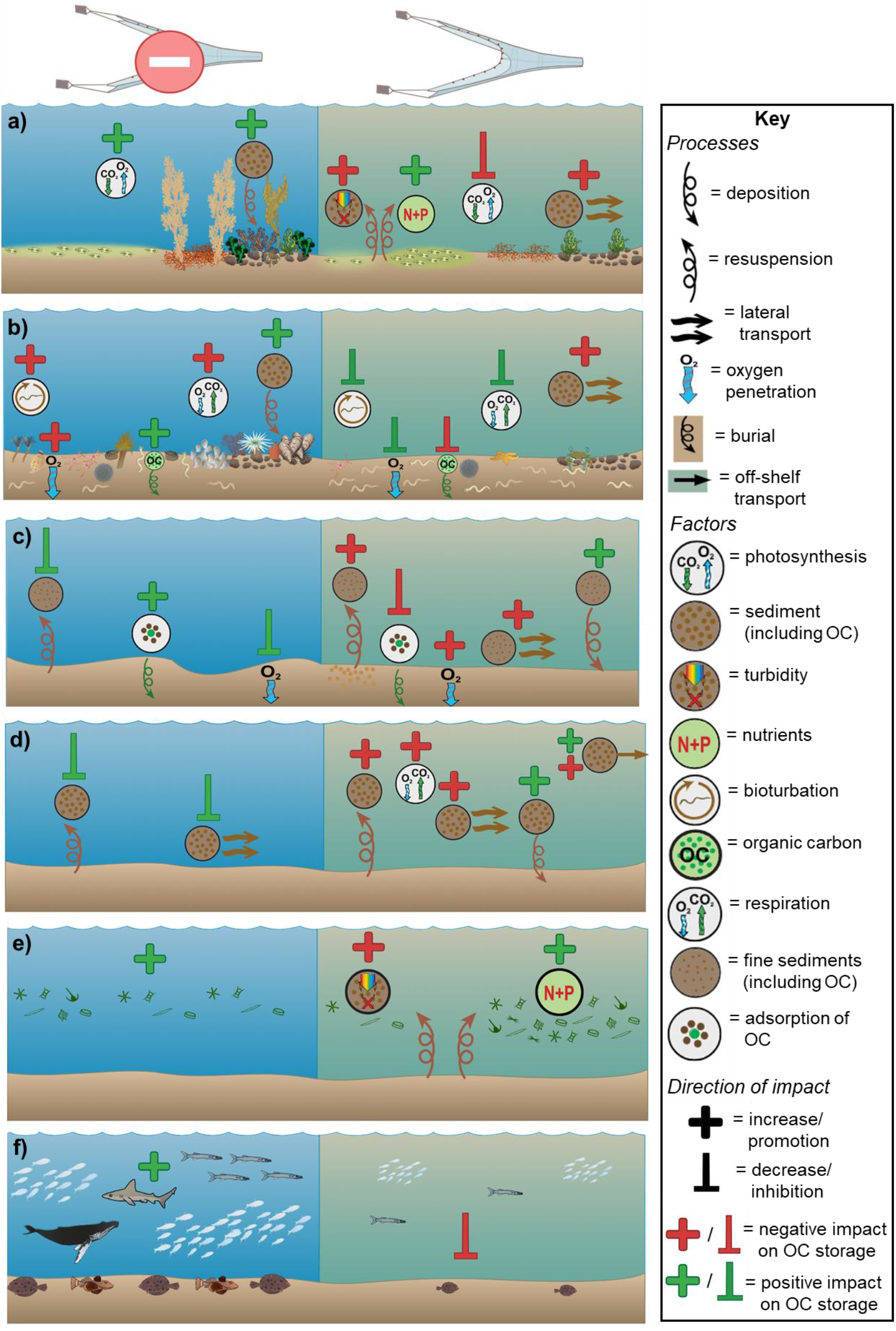
The effects of mobile demersal fishing activity (right) and absence of demersal fishing activity (left) on a) benthic algae, b) benthic infauna and epifauna, c) sediment characteristics, d) sediment dynamics, e) pelagic primary production, f) vertebrate fauna, and how each of these changes may impact seabed sediment organic carbon (OC) stores. Addition symbols indicate when a factor/process would be expected to increase in the presence/absence of fishing; inhibitory arrows indicate when a factor/process would be expected to decrease. The colour of the addition/inhibition symbols indicates whether this change is predicted to impact OC sequestration and storage either positively (green) or negatively (red). Symbols courtesy of Integration and Application Network (ian.umces.edu/media-library)

In some cases, especially in oligotrophic environments, disturbance from mobile demersal fishing may release nutrients from sub-surface sediments and promote primary production, increasing the density of microphytobenthos (Fig. 1a) (Fanning et al. 1982, Falcão et al. 2003, Dounas et al. 2007, Palanques et al. 2014). Counteracting this, sediment suspended by fishing increases turbidity (Ruffin 1998, Palanques et al. 2001) which reduces light penetration and thus photosynthetic rates (Fig. 1a) (MacIntyre et al. 1996). For example, there are contrasting results from fishing impact studies within the Northeast Atlantic. Experimental trawling activity in the Frisian Front significantly reduced chlorophyll a concentration at the sediment surface (Tiano et al. 2019), whereas investigations over a range of trawling intensities in muddy sediments of the Irish Sea found a positive correlation between chlorophyll a concentration and fishing frequency (Sciberras et al. 2016).

In most settings, high frequency mobile demersal fishing would be expected to reduce the abundance of benthic flora on euphotic sediments and to therefore limit the quantity of OC stored directly, and via secondary production (Fig. 1a) (Miller et al. 1996, Middelburg 2018, Mandal et al. 2021). Additionally, benthic micro- and macroalgae are known to increase the stability and accumulation rate of seabed sediments (Yallop et al. 1994, Miller et al. 1996), a primary driver of OC burial and storage (Middelburg 2018, LaRowe et al. 2020). This represents a further mechanism through which disturbance of benthic algae from mobile demersal fishing would limit the potential sequestration rate of OC within sedimentary seabed habitats (Fig. 1a).

### 2.2 Benthic faunal production and processing of OC

The impact of mobile demersal fishing gears on benthic fauna has been widely studied. Long-term effects on community structure and faunal biomass are site-specific and fishing gear dependent (Collie et al. 2000, Kaiser et al. 2002, Thrush and Dayton 2002, Kaiser et al. 2006, Hiddink et al. 2017, Sciberras et al. 2018). Gears which penetrate most deeply into sediment, such as dredges and hydraulic gears, tend to have greater impact than gears with less penetration, such as demersal seines and otter trawls (Collie et al. 2000, Kaiser et al. 2006, Hiddink et al. 2017, Sciberras et al. 2018), although habitat type also has an influence (Rijnsdorp et al. 2020). The largest impacts follow initial experimental trawling events or are seen when comparisons are made to an area of long standing protection (Thrush and Dayton 2002, Cook et al. 2013). Many studies may underestimate the damage done by mobile fishing gears and overestimate speed of recovery because they measure recovery of areas already impacted (Collie et al. 2000, Kaiser et al. 2002, Kaiser et al. 2006, Hinz et al. 2009, Cook et al. 2013, Hiddink et al. 2017, Sciberras et al. 2018).

To a greater or lesser extent, bottom trawling and dredging reduce total benthic biomass and production of benthic fauna and cause loss in abundance and diversity of sessile epifauna and long-lived shallow burrowing infauna (Kaiser et al. 2002, Queirós et al. 2006, Tillin et al. 2006, Sciberras et al. 2018, Tiano et al. 2020). Long-term fishing with mobile gears leads to preponderance of small-bodied, opportunistic, motile infauna, and larger, highly vagrant, scavenging macrofauna (Kaiser et al. 2002, Thrush and Dayton 2002, Kaiser et al. 2006, Tillin et al. 2006). But even within the largely resistant opportunistic meiofauna, mobile demersal fishing still affects diversity and community structure (Schratzberger et al. 2009, Pusceddu et al. 2014).

Benthic fauna are primary drivers of carbon cycling in sediments (Middelburg 2018, Snelgrove et al. 2018, LaRowe et al. 2020, Rühl et al. 2020). For example, in a well-studied area off Vancouver Island, taxonomic and functional richness of benthic fauna explained a similar proportion of variance in pelagic-benthic nutrient flux (~20%) when compared to a suite of environmental variables (Belley and Snelgrove 2016, 2017). Much of the OC that reaches the seabed is directly consumed by deposit and suspension feeding fauna, and is thereafter incorporated into biomass, expelled as faeces and pseudofaeces, or metabolised and remineralised through respiration (Arndt et al. 2013, Keil 2017, Middelburg 2018, Snelgrove et al. 2018, Rühl et al. 2020). While respiration reduces the concentration of OC available for burial and storage, the consumption and processing of OC by benthic fauna may increase proportions of refractory compounds resistant to microbial decomposition, or form specific OC-mineral interactions which can isolate the OC from remineralisation processes, thus improving the potential for burial and long-term storage (Fig. 1b) (Arndt et al. 2013, Middelburg 2018, LaRowe et al. 2020).

Bioturbation and bio-irrigation activities generally increase OC remineralisation due to oxygenation of surface sediments and an increase in the concentration of other electron acceptors, therefore promoting microbial degradation (Fig. 1b) (Hulthe et al. 1998, Arndt et al. 2013, Keil 2017, Snelgrove et al. 2018, LaRowe et al. 2020). However, these activities also transport OC rich surface sediments to deeper sediment layers, potentially increasing their chance of burial and long-term storage (Fig. 1b) (van der Molen et al. 2012, Middelburg 2018, Snelgrove et al. 2018, Rühl et al. 2020, De Borger et al. 2021). For example, in the North Sea, bioturbation by infauna has been found to promote remineralisation by exposing buried material to oxygen (Hulthe et al. 1998) while also being significant in moving carbon from the surface to deeper sediment layers (van der Molen et al. 2012, Middelburg 2018).

The composition and abundance of benthic fauna can also influence the stability and accumulation rates of sediment, which are key drivers of OC burial and storage (Middelburg 2018, LaRowe et al. 2020). While increased bioturbation activity generally has a destabilising effect, burrowing fauna can increase the stability and accumulation rate of sediment if there is an increase in biogenic material such as worm tubes or mucus production, or an increase in structural complexity at the sediment surface from the presence of sedentary and sessile epifauna and biogenic habitat (Fig. 1b) (Ekdale et al. 1984, Thrush and Dayton 2002, Roberts 2007, Borsje et al. 2014, Rühl et al. 2020). For example, in fine sands and muds of the Northeast Atlantic the presence of the tube building polychaete *Lanice conchilega* can lead to increased sediment accretion rates due to changes in flow dynamics around the worm tubes, with impacts on sedimentation dynamics beyond the biogenic structure and over a longer duration than the lifetime of the individual worm (Borsje et al. 2014). In contrast, in the same habitat the density of the Manila clam (*Ruditapes philippinarum*) was positively correlated to sediment erosion rates due to enhanced bioturbation activities (Sgro et al. 2005).

Faunal biomass and production are some of the main contributors of OC in seabed sediments. Therefore, the expected overall impact of mobile demersal fishing on faunal mediated processes is a reduction in OC storage, with the effect somewhat offset by reduced bioturbation and respiration causing lower remineralisation rates. Where the balance lies depends on the many complex interactions discussed above, which are site-specific.

### 2.3 Alteration to sediment composition

Mobile demersal fishing gears can alter the granulometry, topography and vertical structuring of seabed sediments (Trimmer et al. 2005, Puig et al. 2012, Martín et al. 2014b, Oberle et al. 2016a, Oberle et al. 2016b), with extent of change influenced by gear used, sediment type, local hydrology and frequency of fishing (Kaiser et al. 2002, Trimmer et al. 2005, Martín et al. 2014b, Oberle et al. 2016b). Gears that penetrate more deeply into sediment and have a larger footprint cause most impact (Kaiser et al. 2002, Martín et al. 2014b, Eigaard et al. 2016). In highly mobile habitats, the structure and composition of sediment may not be greatly altered by mobile demersal fishing due to strong natural forcing mechanisms, while those found in less hydrologically active environments could be highly affected (Kaiser et al. 2002, Trimmer et al. 2005, Martín et al. 2014b, Oberle et al. 2016b). However, greater sediment mobility may itself be a consequence of long-term use of mobile fishing gears, due to loss of fauna and flora that stabilise sediments (Roberts 2007).

Topographic alterations from mobile fishing gears can consist of visible trawl/dredge tracks and homogenisation in large-scale seabed topography (Kaiser et al. 2002, Martín et al. 2014b, Palanques et al. 2014, Eigaard et al. 2016, Oberle et al. 2016a, Oberle et al. 2016b, Tiano et al. 2020). For example, multibeam surveys have shown that chronic trawling on the continental slopes of the Palamós canyon in the Northwest Mediterranean has had drastic flattening effects on soft sediments (Puig et al. 2012). Mobile demersal fishing also mixes and overturns the top layer of seabed, generally causing a homogenisation of the sediment structure and an increase in density of surface sediments (Martín et al. 2014a, Pusceddu et al. 2014, Oberle et al. 2016b, Paradis et al. 2019). The sediment’s vertical profile can also be altered, with an increase in coarse material towards the surface, caused by winnowing, resuspension and loss of fine material (Fig. 1c) (Martín et al. 2014a, Martín et al. 2014b, Palanques et al. 2014, Pusceddu et al. 2014, Mengual et al. 2016, Oberle et al. 2016b, Paradis et al. 2019). If the local hydrology is relatively depositional, the sediment may be overlain by a surface layer of fine material from the re-deposition of fine sediment which has been re-suspended from deeper layers (Palanques et al. 2014, Oberle et al. 2016b, Tiano et al. 2020). On the Northwest Iberian shelf, all these processes and impacts were identified within a study across different trawling intensities and environmental settings, highlighting the complexity in predicting fine-scale effects of mobile demersal fishing on sediment structure (Oberle et al. 2016b).

The loss of fine, flocculent material and OC-mineral interactions via mobile demersal fishing is another mechanism by which OC sequestration could be reduced (Fig. 1c) (Martín et al. 2014b, Pusceddu et al. 2014, Oberle et al. 2016a). The physical mixing of surface sediments generally causes an increase in oxygen penetration (Martín et al. 2014a, Tiano et al. 2019, De Borger et al. 2021), resulting in reduced OC-mineral interactions (Arnarson and Keil 2007, Estes et al. 2019) and increased microbial respiration and remineralisation (Fig. 1c) (Kristensen et al. 1995, Dauwe et al. 2001, Keil 2017, van de Velde et al. 2018). The process of physical encapsulation of OC by sediment particles and the resultant protection from remineralisation, is seen as a key process in long term OC storage (Burdige 2007, Arndt et al. 2013, Estes et al. 2019, LaRowe et al. 2020). For example, in sediment samples from the Northeast Pacific coasts of Mexico and Washington state, 50% of the oldest OC stores were sorbed to mineral surfaces (Arnarson and Keil 2007).

Further, due to their often biological origin, fine grained sediments such as silts and clays typically have higher concentrations of OC compared to habitats dominated by sand and coarse sediment (Burdige 2007, Paradis et al. 2021, Smeaton et al. 2021). As mobile demersal fishing generally exposes or suspends fine material, this would tend to reduce overall OC storage through resuspension, oxidation and remineralisation (Fig. 1c). Finally, mobile demersal fishing can lead to “organic matter priming”, whereby more easily degraded “labile” OC at the surface is mixed with less easily degraded “recalcitrant” material. This can lead to significantly increased total OC remineralisation rates, although the process is known to vary between environmental settings (van Nugteren et al. 2009, Bengtsson et al. 2018, Riekenberg et al. 2020).

### 2.4 Sediment resuspension and transport

Large volumes of seabed sediments are sufficiently dynamic to be moved laterally and vertically, and become resuspended in the water column by tides, waves and storms (Soulsby 1997, Winterwerp and Kranenburg 2002, Ferré et al. 2008). Mobile demersal fishing activities have at times been shown to exceed, or be a large contributor to, the quantities of sediment displaced by natural forcing mechanisms (Jones 1992, Pusceddu et al. 2005b, Ferré et al. 2008, Martín et al. 2014b, Pusceddu et al. 2015, Mengual et al. 2016, Oberle et al. 2016a, Paradis et al. 2018). Magnitudes involved are highly dependent on depth, gear and sediment type, with deeper penetrating gears and finer sediments causing larger dispersed volumes (Churchill 1989, Ruffin 1998, Durrieu de Madron et al. 2005, Pusceddu et al. 2005b, Ferré et al. 2008, O’Neill and Summerbell 2011, Martín et al. 2014b, Palanques et al. 2014, Mengual et al. 2016, Oberle et al. 2016a). Depending on local hydrographic conditions, sediment may remain in suspension for extended periods of time, and can be transported across large vertical and lateral distances (Durrieu de Madron et al. 2005, Martín et al. 2006, Palanques et al. 2006, Ferré et al. 2008, Martín et al. 2008, Martín et al. 2014b, Palanques et al. 2014, Pusceddu et al. 2015, Oberle et al. 2016a). In the Northern Mediterranean, otter trawling resulted in average suspended sediment concentrations ranging between 6 - 50 mg/l, depending on the study site (Palanques et al. 2001, Durrieu de Madron et al. 2005). The sediment within the water column was found to persist for up to five days (Palanques et al. 2001), while off-shelf transport was 1.4 - 9 times higher when compared to sediment volumes without trawling (Ferré et al. 2008, Palanques et al. 2014). The loss of seabed topography, as discussed above (Puig et al. 2012, Martín et al. 2014b, Oberle et al. 2016b), may also alter local-scale hydrographic conditions, increasing sediment boundary water flows and the magnitude of sediment resuspension (Smith and McLean 1977, Soulsby 1997).

Natural sediment disturbance during storms is known to stimulate increased water column microbial production (Cotner et al. 2000) and higher OC remineralisation rates (Wainright and Hopkinson Jr 1997, Pusceddu et al. 2005b). In general, the resuspension and transport of sediment from mobile demersal fishing will lead to a reduction in OC content (Pusceddu et al. 2005b, Martín et al. 2006, Pusceddu et al. 2015), largely due to increased oxygen exposure times and shifts between anoxic and oxic states, which generally increase remineralisation rates (Fig. 1d) (Kristensen et al. 1995, Hulthe et al. 1998, Dauwe et al. 2001, Keil 2017). Aerobic remineralisation in marine sediments has been measured at between four and ten times faster than in anaerobic conditions, however this is known to vary depending on environmental settings (Kristensen et al. 1995, Hulthe et al. 1998). Fishing induced disturbance may further promote remineralisation, as sediment which is deposited under oxic conditions, buried under anoxia and re-exposed to oxygen can promote especially high OC degradation rates (Hulthe et al. 1998). This has been identified in the biochemical signature of suspended particulate OC within trawling grounds of the North Mediterranean, with a significant shift from labile to refractory OC compounds (Pusceddu et al. 2005a, Pusceddu et al. 2005b, Pusceddu et al. 2015).

Previous studies have shown that it is challenging to fully quantify the amount of OC that will be remineralised after disturbance, rather than simply being moved elsewhere (Wainright and Hopkinson Jr 1997, Pusceddu et al. 2005b, Martín et al. 2006, Martín et al. 2008, Lovelock et al. 2017). There is also the potential that sediment resuspension from mobile demersal fishing could increase OC storage in adjacent areas (Fig. 1d). This could occur from higher sedimentation rates near to fishing grounds leading to increased burial of OC which is already present within the seabed, or burial of benthic algae and sessile fauna (Churchill 1989, Jones 1992, O’Neill and Summerbell 2011, Oberle et al. 2016a, Sciberras et al. 2016). It could also lead to transportation of OC-rich shelf and slope sediments (Atwood et al. 2020) to deeper waters below mixing depths (Fig. 1d) (Caldeira et al. 2002, Martín et al. 2006, Ferré et al. 2008, Martín et al. 2008, Paradis et al. 2018, Legge et al. 2020). Such off-shelf induced transport of sediment and OC has been recorded as deep as 1750 m in continental slope trawling grounds of the Palamós canyon in the Northwest Mediterranean (Martín et al. 2006, Palanques et al. 2006, Martín et al. 2008).

Overall, increased sediment resuspension from mobile demersal fishing would be expected to reduce the current store of OC in seabed sediments due to disturbance of accumulations and increased oxygen exposure times (Keil 2017, Luisetti et al. 2019, De Borger et al. 2021). Future sequestration would also be limited as newly settling organic material would be kept in suspension, precluding it from burial and storage (Churchill 1989, Ruffin 1998, Martín et al. 2014b, Oberle et al. 2016a). The magnitude of impact, however, will be largely based on local hydrography, which primarily determines the fate of resuspended OC (Wainright and Hopkinson Jr 1997, Ferré et al. 2008).

### 2.5 Alteration in pelagic primary production

As most seabed OC is allochthonous, the total amount which reaches seabed sediments is strongly driven by the level of primary production in the overlying water column (Seiter et al. 2004, Turner 2015, Atwood et al. 2020). As noted previously, sediment disturbance by mobile fishing gears, or natural forces, can release significant concentrations of nutrients into the water column (Fanning et al. 1982, Falcão et al. 2003, Polymenakou et al. 2005, Palanques et al. 2014). In shallower areas, released nutrients will likely enter into or remain in the euphotic zone, where their fertilisation effect can increase phytoplankton primary production (Fig. 1e), (Fanning et al. 1982, Dounas et al. 2007, Palanques et al. 2014). For example, modelling predictions from trawling experiments in the Eastern Mediterranean at Heraklion Bay, estimate that nutrient upwelling from bottom trawling could increase net annual primary production by 15% (Dounas et al. 2007) with subsequent settlement raising OC in seabed sediments (Falcão et al. 2003, Polymenakou et al. 2005, Palanques et al. 2014, Turner 2015). Alongside this, as discussed for microphytobenthos, demersal fishing activity can also reduce rates of photosynthesis by increasing turbidity (Fig. 1e), (Ruffin 1998, Palanques et al. 2001, Adriano et al. 2005, Cloern et al. 2014).

### 2.6 The contribution of vertebrate fauna to OC storage

Although not a focus of this review, the removal of vertebrate species by benthic and pelagic fisheries could influence the mass of OC stored in seabed sediments (Pershing et al. 2010, Atwood et al. 2015, Mariani et al. 2020). The emerging field of “fish carbon” describes the contribution of vertebrate fauna to OC storage and sequestration within seabed sediments from defecation, pelagic mixing, bioturbation, trophic interactions and deadfall (Trueman et al. 2014, Turner 2015, Saba et al. 2021). Although the magnitudes of effect are poorly resolved, the reduction in population size and average body size of marine vertebrates that results from over-harvest, is expected to reduce the amount of carbon exported to the seabed (Fig. 1f) (Pershing et al. 2010, Trueman et al. 2014, Atwood et al. 2015, Mariani et al. 2020). For example, since 1950, the combined catch of Tuna, Mackerel, Shark and Billfish is estimated to have prevented approximately 21.8 Mt of OC being stored in seabed sediments (Mariani et al. 2020). The removal of predatory vertebrates will also cause trophic cascades, potentially leading to alterations in benthic faunal communities, triggering the feedback mechanisms on OC discussed above (Atwood et al. 2015).

### 2.7 Interactions and feedback mechanisms

All factors discussed here interact in a variety of positive and negative feedback loops. For example, alterations in sediment characteristics will influence the community structure of benthic flora and fauna, and vice versa. Additionally, pelagic primary production, trophic interactions, and the abundance and community composition of vertebrate fauna will all further alter benthic population changes induced by mobile demersal fishing. These factors are also influenced by chemical and physical oceanographic processes that are outside the scope of this review. The structure and diversity of the microbiome is also strongly influenced by the composition of benthic flora and fauna (Middelburg 2018, LaRowe et al. 2020, Rühl et al. 2020). However, the microbiome itself can be impacted by mobile demersal fishing activities adding further complexity to the overall picture (Watling et al. 2001, Polymenakou et al. 2005).

## 3. Experimental results

From a systematic literature search (see Supplementary material), 40 peer-reviewed studies were identified which investigated the impact of mobile demersal fishing on the seabed, and directly measured OC or organic matter (OM) and/or remineralisation rates in seabed sediments (Table 1). The 40 studies covered 12 oceanic realms with greatest representation from the Northeast Atlantic (43%), Mediterranean (23%) and Northwest Atlantic (15%). The majority of studies (58%) investigated the impacts of commercial fishing activities. The remainder either used experimental trawling/dredging methods (33%), a mixture of experimental trawling and monitoring of commercial fishing (5%), or mathematical modelling of fishing impacts (5%). A variety of experimental setups were employed including impact-control site comparisons (43%), before-after fishing impact (23%), and low-high impact contrasts which lacked controls (20%). Additionally, 13% of studies used a before-after control-impact design either alone or in combination with an impact-control experiment; and one investigated the recovery of seabed sediment OC after a long-term closure to mobile demersal fishing (Table 1). It should be noted that for many of these studies, in areas considered “control sites” there is the potential for them to still be affected by mobile demersal fishing activities. This often occurs due to insufficient monitoring (e.g. no Vessel Monitoring System data on smaller vessels), lack of enforcement (i.e. within a supposed closed area) or lack of recovery time since cessation of fishing, particularly given the long timescales of recovery expected for many habitats (Roberts 2007).

**Table 1.**
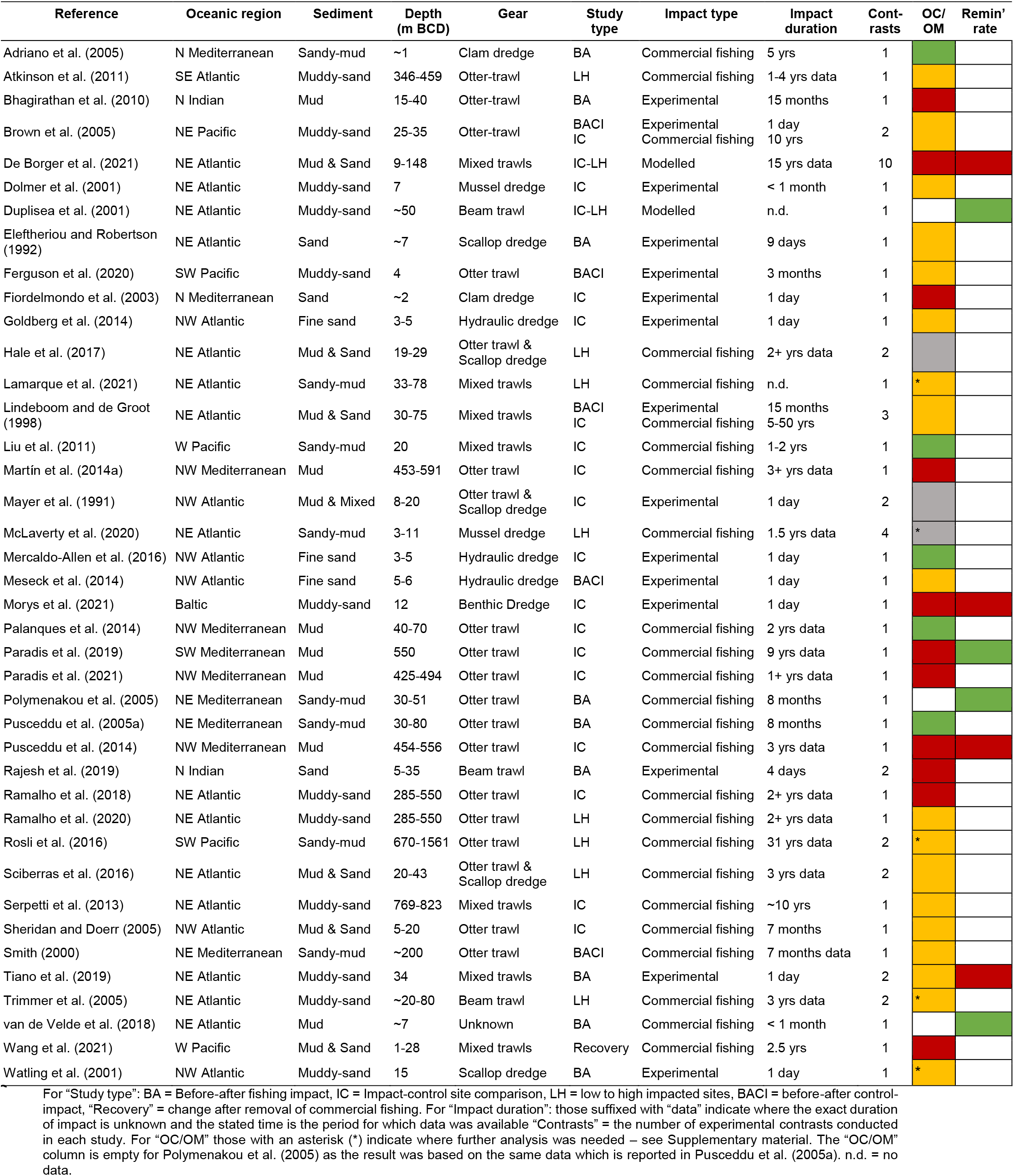
Summary of studies which investigated the impact of mobile demersal fishing on the seabed and directly measured organic carbon (OC) or organic matter (OM), and/or remineralisation rates of OC/OM in the sediment. The last two columns indicate whether the presence or increase in demersal fishing activity was reported to cause lower (red), higher (green), no significant effect (orange) or mixed effects (grey) in the concentration or mass of OC/OM (“OC/OM”), or organic carbon remineralisation rate (“Remin’ rate”), within seabed sediments.

Of the 40 studies identified, 11 investigated the effect of mobile demersal fishing across multiple sites, habitat types or gear-types, and made inferences for each investigation separately (Table 1); this produced a total of 62 experimental contrasts (Table S1). Of these, 59 measured changes in OC/OM concentration. A finding of no significant effect was reported in 51% of contrasts; 41% reported lower OC in fished sites compared to unfished control sites (or in areas with higher fishing intensities), with 8% reporting higher OC (Table S1).

Studies which reported a negative impact from mobile demersal fishing on OC generally occurred in muddy sediments, while those which reported higher OC in response to this disturbance, or no effect, occurred in a mixture of sandy and muddy sediments (Table S1). On average, the duration of impact was higher for studies which reported a negative effect of demersal fishing on OC, when compared to those which reported a positive or non-significant effect, with estimated values of median impact duration at 36 months and 18 months respectively (Table S1). Most that reported a negative impact from demersal fishing were Impact-Control studies (75%) or Before-After fishing impact studies (13%). In contrast, those that reported no significant effects were predominantly Low-High impact studies lacking controls (43%) and Impact-Control studies (27%). The 5 studies which reported an increase in OC were relatively evenly spread between Impact-Control designs (60%) and Before-After designs (40%). The median depth at which the research was conducted was relatively similar between different experimental outcomes, with median depths of 22 m, 31 m and 20 m, for studies which reported a decrease, no significant effect, and an increase in OC respectively (Table S1).

Within the literature examined, there were 18 inferences about the impact of mobile demersal fishing pressure on sediment carbon remineralisation rate. Of these, 78% reported that demersal fishing activity decreased remineralisation rate in seabed sediments, with the rest concluding opposite (Table S1). Although no clear trend was identified between studies, it seems the result is highly dependent on local hydrographic conditions. For example, in more depositional environments, mobile demersal fishing may cause oxygenation of sediments and redeposition of recently expulsed organic material back to the seabed, leading to an increase in remineralisation rate (Duplisea et al. 2001, Polymenakou et al. 2005, van de Velde et al. 2018). In more hydrologically active environments, resuspension and lateral/vertical transport of sediments would be expected to reduce OC in surface sediments which, along with removal of fauna, could limit the rate of remineralisation (Pusceddu et al. 2014, Tiano et al. 2019, De Borger et al. 2021, Morys et al. 2021).

The evidence discussed earlier in Section 2 would suggest that removing or reducing demersal fishing pressure from the seabed would have net benefits to carbon sequestration and storage. However, the experimental results identified here indicate study-specific and site-specific outcomes. The majority of studies which identified no significant effect in sediment OC used an experimental design which compared sites with different magnitudes of fishing impact but lacked controls. There is a clear need for further identification and investigation of seabed sediment habitats that have had true long-term protection from demersal fishing. Those studies which reported an increase in OC, or no effect, also generally occurred in sandier sediments which may be subjected to higher levels of natural disturbance; however, as highlighted in this review, there will also be sandier areas where the impact of fishing activity outweighs natural forcing mechanisms. Finally, many of the studies identified in the systematic review were not primarily designed to investigate the impact of demersal fishing on carbon storage or remineralisation, although this may not affect the direction of their conclusions.

## 4. Future research

As highlighted by the varied results, there is a clear need for further research into the potential impact of mobile demersal fishing on OC sequestration and storage in seabed sediments. Recent first order estimates have suggested that globally, mobile demersal fishing could remineralise between 160 - 400 Mt of OC from marine sediment stores annually (Sala et al. 2021). It has also been suggested that historical trawling on global continental slopes could have removed ~6000 Mt of OC from the upper-most centimetre of sediment alone (Paradis et al. 2021). In addition, it has been estimated that ~2 Mt of OC is remineralised from UK shelf sediments each year by mobile demersal fishing (Luisetti et al. 2019). Although these estimates contain large generalisations, their scale reveals the massive potential for mobile demersal fishing to reduce carbon stores.

Following disturbance by mobile demersal fishing a proportion of OC will be remineralised in the benthos or in the water column, however some will simply remain in-situ and be re-buried, and a further proportion will be transported over a range of distances either being consumed or re-buried (Pendleton et al. 2012, Lovelock et al. 2017). A key research gap is the quantification of OC that follows each of these processes in different environmental settings and under different types of fishing impact. Sala et al. (2021) only account for remineralisation of disturbed OC which remains in-situ or resettles within 1 km^2^, as they consider the fate of sediment which stays in suspension as unknown. In their paper, Sala et al. (2021) consider that 87% of the OC disturbed remains in-situ or resettles uniformly across global fishing effort, and of this anything between 1-69.3% will be remineralised, with the magnitude dependent upon two relatively coarse metrics, namely: estimated proportion of OC which is labile, and oceanic basin degradation rate. In contrast, Luisetti et al. (2019) use an upper estimate that 100% of the OC resuspended by mobile demersal fishing will be remineralised, but they do not consider the fate of OC that is disturbed but remains in-situ. Although both studies give a representation to the scale of OC which may be lost, improved quantification of these metrics is clearly needed before accurate measures of OC lost, or inorganic carbon produced, can be quantified. OC in seabed sediments is not naturally inert, passing through a range of aerobic and anaerobic remineralisation pathways to varying sediment depths. Thus more consideration is needed to understand the influence of natural remineralisation rates within seabed sediments under different environmental settings, and therefore quantify the additional effect of mobile demersal fishing in each area. In seabed sediment habitats with high hydrodynamic activity, low deposition rates, and high oxygen penetration depths, the additional disturbance of demersal fishing on OC may be more limited.

We must also consider the cumulative or finite nature of disturbance by demersal mobile fishing on OC stores. It is not clear how much of the estimated 360 Gt of OC in the top 1 m of sediment is actually under threat (Atwood et al. 2020). While mobile demersal fishing can only penetrate between around 2 and 20 cm into the sediment (Hiddink et al. 2017), repeated chronic impacts may continue to disturb and displace sediment more deeply (Sala et al. 2021). It is possible, that in chronically fished areas significant further loss of OC stores will not occur due to historic depletion in OC stocks. However, in such areas carbon sequestration and accumulation of OC would be limited by the frequency of disturbance to newly settled material (Sala et al. 2021). By contrast, if new fishing grounds emerge, these could act as huge sources of carbon emissions as sediment becomes disturbed and OC is remineralised (Gogarty et al. 2020).

There is also a need to identify a clear baseline from which changes in OC can be measured. Standing stock of OC in global seabed sediments is relatively well resolved at a number of spatial scales (e.g. Seiter et al. 2004, Lee et al. 2019, Luisetti et al. 2019, Atwood et al. 2020, Legge et al. 2020, Diesing et al. 2021, Smeaton et al. 2021). However, precise estimates of OC remineralisation, accumulation and burial rates are generally lacking (Berner 1982, Burdige 2007, Keil 2017, Wilkinson et al. 2018, Luisetti et al. 2019, Legge et al. 2020, Diesing et al. 2021). Any studies which aim to quantify the impact of demersal fishing on carbon storage and sequestration must therefore quantify both the before and after scenarios for robust conclusions to be drawn.

It is important that future research into the impact of mobile demersal fishing on carbon storage is focused in areas which are expected to contain significant stocks of OC or have large future sequestration potential, based on their geographic projections (Atwood et al. 2020), sediment characteristics (Smeaton et al. 2021) and local hydrology (Lee et al. 2019). Research should also focus on areas that overlap with significant mobile demersal fishing pressure (Amoroso et al. 2018, Kroodsma et al. 2018, Sala et al. 2021), and where this can be compared to areas that could be considered truly “unfished”, either from well enforced protected areas or specific environmental settings.

On land, retrospective analyses of changes in human use and vegetation cover have been critical to estimating how people have altered the planetary carbon cycle. It is vital that this historical context is considered when further investigating the potential impact of mobile demersal fishing on global seabed OC sequestration and storage, and the opportunities for recovery if this pressure is removed. Due to the extended timeframes needed for some seabed habitats to fully recover, true long-term protection and monitoring of OC is needed to fully deduce carbon storage potential. Without considering areas of seabed that have experienced genuine long-term protection, it is not possible to gain a true baseline from which impacts can be compared (Pinnegar and Engelhard 2008). Within this review, only one study could be found which looked at the direct recovery of OC in seabed sediments following the medium-to-long term removal of fishing pressure (Wang et al. 2021). Gaining further evidence is vital to understand how much OC can accumulate when mobile demersal fishing is removed, and how this may change over the course of recovery.

## 5. Concluding remarks

Seabed sediments are one of the planet’s primary OC stores and strongly influence the oceanic sink for atmospheric CO_2_ (Gruber et al. 2019, Atwood et al. 2020, Watson et al. 2020, Sala et al. 2021). It is an urgent priority to better understand the effect of mobile fishing gear use on seabed OC sequestration and storage, and to incorporate clear blue carbon considerations into global seabed management. As only around 2-3% of the world’s seabed is currently closed to trawling and dredging (Roberts et al. 2017, Marine Conservation Institute 2021), increasing the scale of protection could offer huge climate change mitigation potential and bring corresponding gains in biodiversity (Roberts et al. 2017, Seddon et al. 2019, Roberts et al. 2020, Sala et al. 2021). Across the world, mobile demersal fisheries are highly fuel inefficient and produce most of the fishing industry’s direct greenhouse gas emissions (Parker et al. 2018). A shift to less damaging fishing methods could provide major net benefits for increasing natural carbon sequestration and storage in the seabed, whilst significantly reducing emissions of CO_2_.

The results of recent regional and global scale publications which calculated first-order estimates of CO_2_ produced from disturbance to seabed sediments by mobile demersal fishing must be taken with both concern and caution (Luisetti et al. 2019, Paradis et al. 2021, Sala et al. 2021). As identified in this review, demersal fishing by trawling and dredging is in many cases likely to limit the storage and sequestration of OC, but to draw firm conclusions more experimental studies covering a wide range of environmental settings, habitat types and fishing pressures is required to address the large number of unknowns and site-specific drivers associated with the status of OC on the seabed.

## Author contributions

GE prepared the original draft and undertook the systematic review. All authors contributed to conceptualisation, writing, reviewing and editing.

