## Supplementary material for "The potential for mobile demersal fishing to reduce carbon storage and sequestration in seabed sediments"

**Details of systematic literature search**

**Article search**

Literature searches were conducted in April 2021 in Web of Science and Scopus. Within Web of Science, its “Core collection” was searched via the field “Topic”, which examines a paper’s title, abstract, author, keywords and “keywords plus”. Within Scopus, the “Advanced search” was run via the field “Title-Abs-Key”, which scans a paper’s title, abstract and keywords. Within both databases the same search terms were used, with results containing at least one term from each of the following groups:

- Sediment* OR Mud* OR Sand* OR Clay* OR Silt* OR Gravel*
- Coast* OR Sea* OR Ocean* OR Estur* OR Estuary OR Marine
- Trawl* OR Dredg* OR “Demersal Fishing” OR “Demersal Fisher*” OR “Bottom Fishing” OR “Bottom Fisher*” OR “Benthic Fishing” OR “Benthic Fisher*”
- “Organic Carbon” OR “Organic Matter” OR “Organic Content” OR “Blue Carbon” OR Remineralisation OR Remineralization OR “Carbon mineralisation” OR “Carbon mineralization”

In addition, a bibliographic search of all relevant articles identified was conducted to ensure all appropriate articles were included. Only English language articles were assessed.

**Inclusion criteria**

To be included in this review studies were required to meet the following characteristics:

- *Habitat -* Any subtidal benthic marine environment, including estuarine; but not typical vegetated blue carbon habitats (saltmarsh, seagrass and mangroves).
- *Fishing pressure* - Included studies had to investigate or quantify demersal mobile fishing pressure (i.e. benthic trawling or dredging for fish or shellfish).
- *Sampling -* Seabed sediments must have been sampled (or modelled) to be included in the review. Studies which only sampled the water column or biota were excluded during screening.
- *Response metric -* Levels of organic carbon (OC), organic matter (OM) or carbon remineralisation rates had to be sampled within the seabed, or used directly as a response variable within each study
- *Interaction -* The presence or magnitude of mobile demersal fishing pressure needed to be related to the response variable of interest in some way. This may have been conducted using a variety of methods such as Before-After impact studies, Impact-Control site comparisons, Before-After removal of pressure, or measures across gradients of pressure. Studies that contained no contrasts in levels of fishing pressure were excluded.

**Screening process**

All articles identified from the searches were exported into a single EndNote library then duplicates removed. Screening was conducted via a hierarchical process that first assessed title, then abstract and finally full text. At each stage an article was assessed against the inclusion criteria described above, with those considered relevant or of unclear relevance passing to the next level of assessment. Once relevant articles were identified, all their bibliographic references were exported to a new EndNote folder by selecting all “Related Records” for each entry in Web of Science. The screening process was then conducted for a second round.

In total, 1,124 articles were identified from the search, of which 367 were duplicates. Removal of the latter lead to 757 screened with 693 excluded at the title and abstract stages. Of the 64 articles assessed at full text, 34 were considered relevant. The reasons why the other 30 were excluded were as follows: one full text was not available; one study consisted of re-analysis and republication of data from a previous study; one study was about seagrass; eight did not consider mobile demersal fishing; four had no measure of seabed sediments; seven did not quantify OC/OM or carbon remineralisation; and eight studies had no contrasts between levels of fishing impact. The second round of screening, which considered all cited literature within the 34 relevant studies, identified a further 4 relevant studies for inclusion. Finally, all authors reviewed the list of identified studies and contributed known relevant papers that were absent or very recently published – adding a further 2 studies.

**Data extraction and analyses**

If individual studies investigated the effect of mobile demersal fishing across multiple sites, habitat types, or gear-types, and made inferences for each investigation separately, these were separated into individual experimental contrasts for discussion within this review. This led to 62 different experimental contrasts identified across the 40 studies examined (Table S1). Qualitative or semi-quantitative descriptive information extracted from each relevant article included the study location, sediment type, water depth, fishing gear type, study design, method for measuring fishing impact, impact duration (or period for which data were available), depth within the sediment to which sampling was conducted, and the frequency of any temporal sampling (see Table S1 for further details). If information on sediment type or the fishing gear was lacking, a grey literature search was conducted into the study-site and/or local fishery to obtain a qualitative description of these characteristics.

In 35 of the 40 studies examined, direct inference on the significance and direction of effects of mobile demersal fishing on OC/OM was made within the publication. This was identified through the display of formal statistical tests or clear statements made within the text. In the five remaining studies, no direct inference was made within the text or supplementary information, however applicable data were presented. In these cases additional analysis was required to infer the impact of demersal mobile fishing on OC/OM (Table S1). In these cases, basic statistical tests were conducted on the data presented within the publication to identify the likely significance and direction of effect. All statistical analyses were carried out in R 4.0.4. Studies which collected data on OC/OM across a continuous gradient of fishing pressure were assessed using Spearman rank correlations, with associated p-values calculated using the *cor.test* function in base R. Those studies which considered OC/OM levels across discrete gradients of pressure were analysed with basic linear models.
